# Convergence of aging- and rejuvenation-related epigenetic alterations on PRC2 targets

**DOI:** 10.1101/2023.06.08.544045

**Authors:** Michael A. Koldobskiy, Oscar Camacho, Pradeep Reddy, Juan Carlos Izpisua Belmonte, Andrew P. Feinberg

## Abstract

Rejuvenation of tissues in physiologically aging mice can be accomplished by long-term partial reprogramming via expression of reprogramming factors (Oct4, Sox2, Klf4 and c-Myc). To investigate the epigenetic determinants of partial reprogramming-mediated rejuvenation, we used whole genome bisulfite sequencing to carry out unbiased comprehensive profiling of DNA methylation changes in skin from mice subjected to partial reprogramming, as well as young and untreated old controls. We found a striking convergence of age- and rejuvenation-related epigenetic alterations on targets of the Polycomb repressive complex 2 (PRC2). These results are also supported by a likewise prominent enrichment of PRC2 targets in gene expression data, suggesting that PRC2 activity can modulate aging and mediate tissue rejuvenation.

## MAIN TEXT

Partial reprogramming using cyclic expression of the transcription factors OCT4, SOX2, KLF4 and c-MYC (OSKM) has emerged as a powerful strategy to reverse age-associated phenotypes^1-5^. Recently, Browder et al. established a long-term partial *in vivo* reprogramming protocol in normal physiologically aging mice, using mice carrying a single copy of an OSKM polycistronic cassette and a reverse tetracycline transactivator (rtTA) in a C57BL/6 genetic background (4F mice)^5^. Analysis of the skin of old treated mice revealed changes consistent with histologic and functional rejuvenation when compared to old untreated mice, including increased epidermal thickness, higher proliferative capacity and decreased fibrosis after injury, reversal of age-related metabolic changes and downregulation of genes involved in inflammation and epidermal differentiation^5^. This suggested that long-term partial reprogramming preserves a more plastic and less differentiated state in aged skin cells.

Given the dramatic reversal of aging-related signatures by long-term partial reprogramming, we sought to understand epigenetic changes that underlie the rejuvenation process. Previously, the application of a DNA methylation array-based aging clock involving fewer than 700 mostly non-functional CpG sites demonstrated a reversal of age-related DNA methylation changes following long-term partial reprogramming^5,6^. Aside from the small number of sites examined, the distinction between biological and chronological components remains a significant challenge of interpretation^7^.

Here we characterize the genome-wide DNA methylation landscape comprehensively in a model of controlled tissue rejuvenation induced by long-term partial reprogramming. We employed whole-genome bisulfite sequencing (WGBS), that can discriminate quantitatively and on a single-read basis methylation status of ∼15 million CpG dinucleotides and is >4 orders of magnitude more than the aging clock CpG site array. We carried out WGBS on skin from five 4F mice treated with long-term partial reprogramming from 15 months of age until 22 months, four old untreated 4F mice (22 months), and three young 4F mice (3 months old) (Table S1). WGBS data was analyzed using informME, a powerful tool for quantifying epigenetic variability by computing DNA methylation potential energy landscapes across the genome, capturing both mean methylation level (MML) and methylation variability as encapsulated by normalized methylation entropy (NME), a version of Shannon entropy that quantifies the disorder of methylation within an analysis region. This method permits identification of genomic regions of significant DNA methylation discordance using the Jensen-Shannon distance (JSD) of information theory, based on differences in probability distributions of methylation rather than conventional differential methylation analysis that is based solely on mean methylation differences^8–11^.

Principal component analysis (PCA) revealed a clear separation between the different groups, capturing the age and rejuvenation-related differences in PC1 and the partial reprogramming treatment-specific effects in PC2 (Fig. 1a). Genome-wide distributions of MML and NME showed a substantial shift towards hypomethylation and a more stochastic and disordered epigenome in old untreated skin when compared to young skin (Fig. 1b,c). Conversely, old treated skin showed an increase in global methylation levels and a reduction of methylation entropy in comparison to old untreated samples, resembling young skin (Fig.1b,d,e). The same trend was consistent when examining MML and NME distributions over selected genomic features (Supplementary Fig. 1).

**Figure 1.**
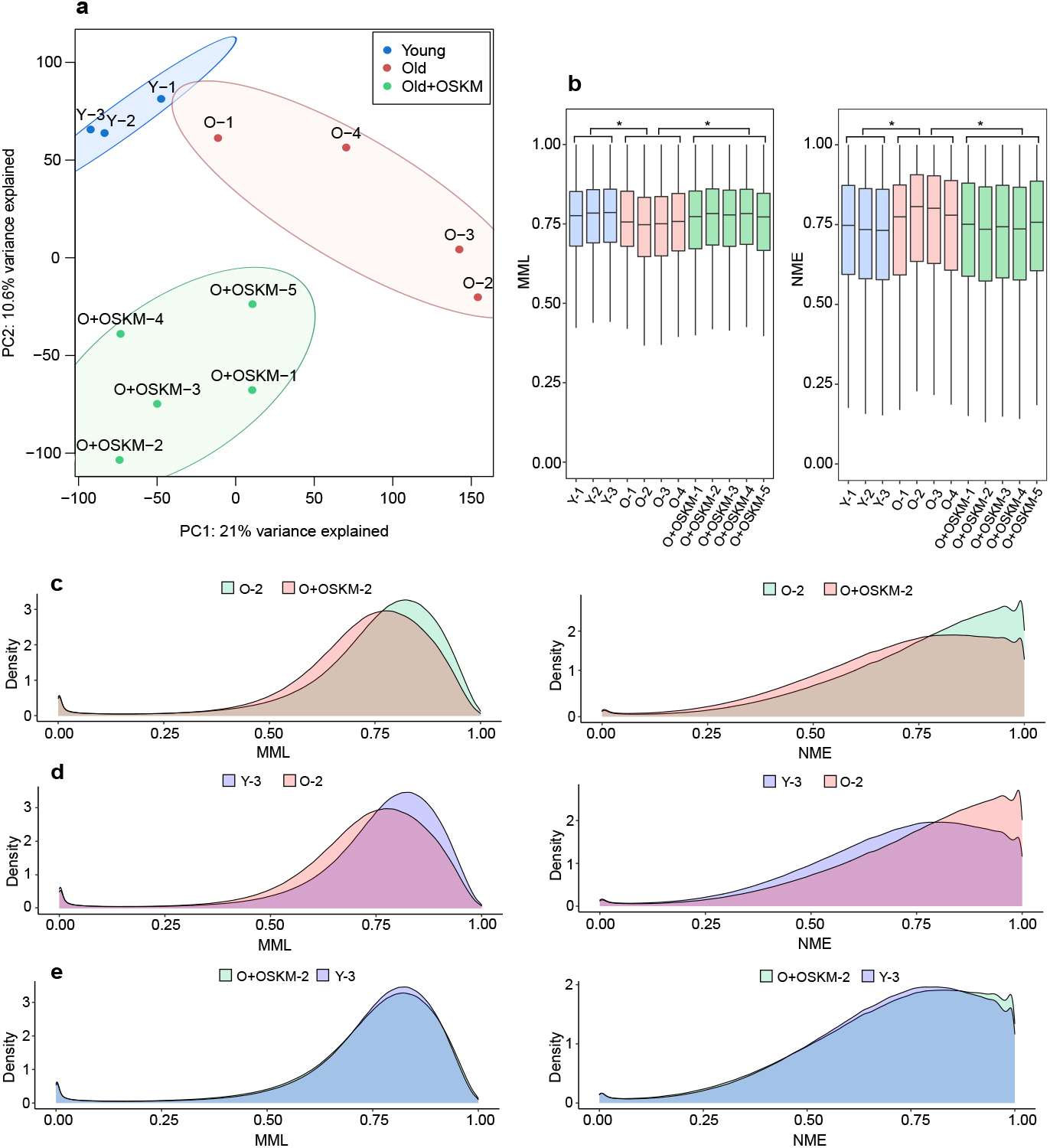
Age-associated genome-wide epigenetic shift is reversed with partial reprogramming-mediated rejuvenation. **A**. Plot of principal components (PC1 and PC2) obtained from PCA of WGBS data corresponding to young, old untreated and old treated (old+OSKM) samples. **B**. Boxplots of genome-wide distributions of mean methylation level (MML) and normalized methylation entropy (NME). **C**. Density plots of genome-wide distributions of MML and NME in representative samples from each group.

Given that aging and long-term partial reprogramming are associated with altered DNA methylation, we next focused on identifying the genomic targets of epigenetic disruption in skin aging and rejuvenation. We ranked genes based on the potential of their DNA methylation states within their promoter and gene body regions to distinguish between young, old untreated and old treated groups, using the mutual information between the methylation state and the phenotype, computed as the average of the sum of squares of all JSD values within analysis regions (Table S2). We identified the most epigenetically discordant genes and evaluated for enrichments among the top 500 genes in each group-wise comparison using Gene Set Enrichment Analysis (GSEA). Remarkably, when comparing both young and old treated to old untreated, we found a high and consistent enrichment in Polycomb repressive complex 2 (PRC2) gene targets (as shown by enrichment in targets of EZH2, SUZ12, EED, JARID2 and MTF2, and genes possessing H3K27me3 marks) (Fig. 2a). The dramatic enrichment observed in both comparisons suggested a convergence of aging- and rejuvenation-related epigenetic changes specifically to genes that are targets of PRC2 activity. When assessing the changes in MML and NME within EZH2 binding sites, regions harboring the H3K27me3 histone mark and poised promoters by employing available ChIP-seq data and chromHMM annotations, we identified a significant shift of the methylation levels towards hypermethylation and a dramatic increase in entropy in the old untreated samples when compared to young skin. Old treated skin exhibited a specific reversal of methylation changes in these genomic regions (Fig. 2b). Interestingly, poised promoters showed the greatest JSD and among the most elevated differences in NME in both young and old treated when compared to old untreated, indicating such poised domains are the most epigenetically dysregulated regions, gain significant entropy with age and such epigenetic disorder can be reversed upon partial reprogramming treatment. Furthermore, as opposed to the global age-related hypomethylation trend in other regions of the genome, poised promoters were the only domains exhibiting an age-related increase in DNA methylation, also reversed with partial reprogramming treatment (Supplementary Fig. 2). An example of a differentially methylated region is shown in Fig. 2c for the cluster *Hoxa7-Hoxa9*, relevant in embryonic development, regeneration and aging.

**Figure 2.**
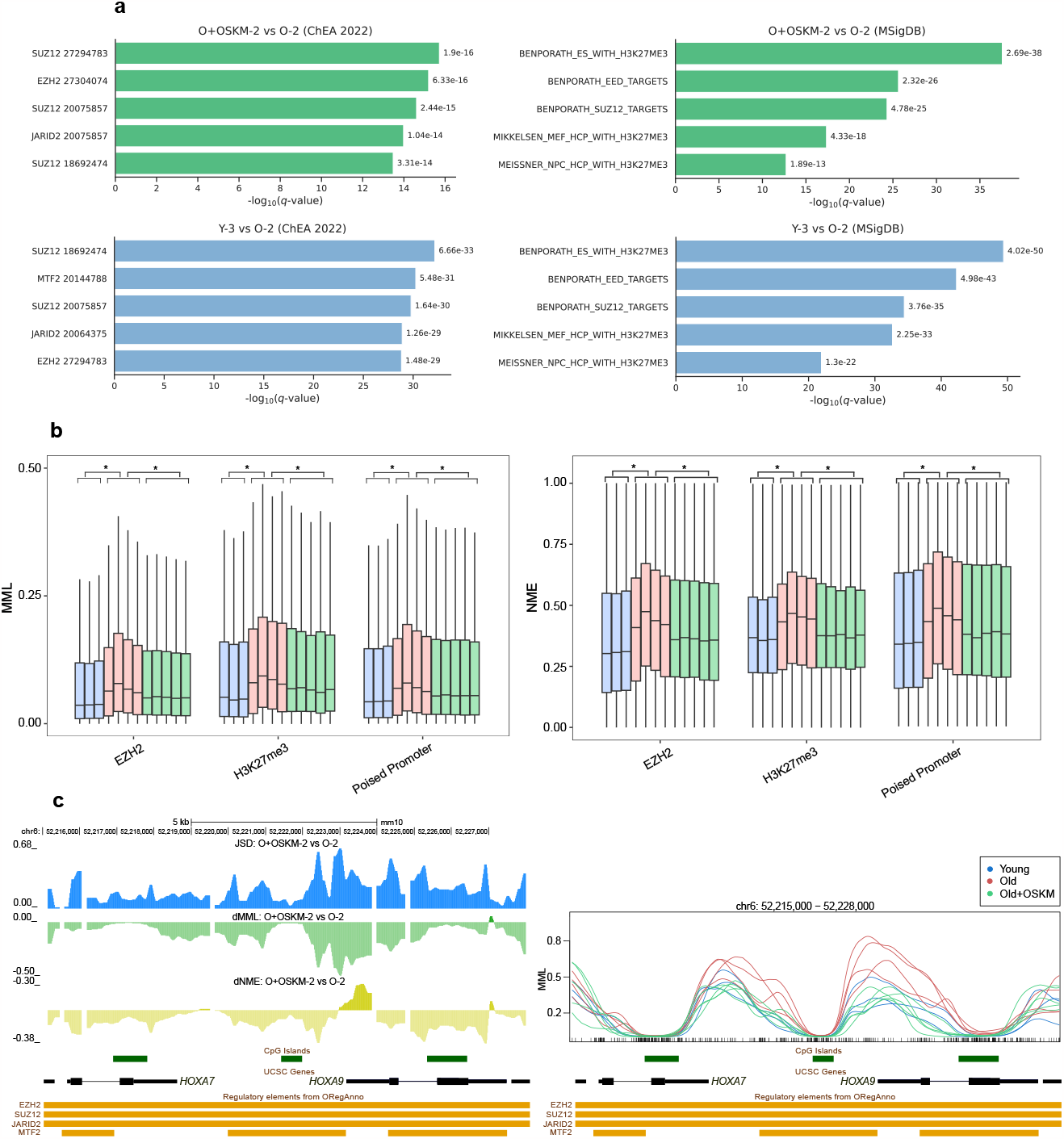
DNA methylation discordance identifies PRC2 targets as major convergence elements in aging and partial reprogramming-mediated rejuvenation. **A**. Enriched gene sets obtained from GSEA of the top 500 epigenetically discordant genes based on JSD in young and old treated (old+OSKM) compared to old untreated comparisons. **B**. Boxplots of MML and NME distributions within EZH2 binding sites, regions harboring H3K27me3 histone mark and poised promoters. **C**. JSD and differences in MML and NME across the cluster *Hoxa7-Hoxa9* in a comparison between representative treated old (O+OSKM-2) and untreated old (O-2) samples.

To evaluate the relationship of these findings to gene expression alterations during aging and partial reprogramming, we carried out differential gene expression (DEG) analysis using RNA-seq data from young, old untreated, and old treated skin samples. Using gene set enrichment analysis (GSEA), we found that among overexpressed DEG in both young and old treated when compared to old untreated skin there was likewise a prominent enrichment of PRC2 targets (Fig. 3), suggesting PRC2 drives age-related gene expression changes. Several genes involved in aging and regeneration found to be both differentially methylated by the partial reprogramming treatment when compared to old untreated and differentially expressed in aging or partial reprogramming, including *Crtc1, Smad3, Wnt3a, Nr4a2, Hmbox1, Foxa1, Camk2b, Egr, Ngf, Pim1, Tbx3, Twist1, Msrb3, Etv6*, and multiple members of the HOX and SOX families, as well as different genes that mediate the age-associated changes in metabolism, remodeling of extracellular matrix and inflammatory response. Among the epigenetically dysregulated genes without apparent change in gene expression, we also found *Foxo3* and *Foxo6*, relevant in aging, and *Jarid2* and *Kdm2b*, epigenetic regulators.

**Figure 3.**
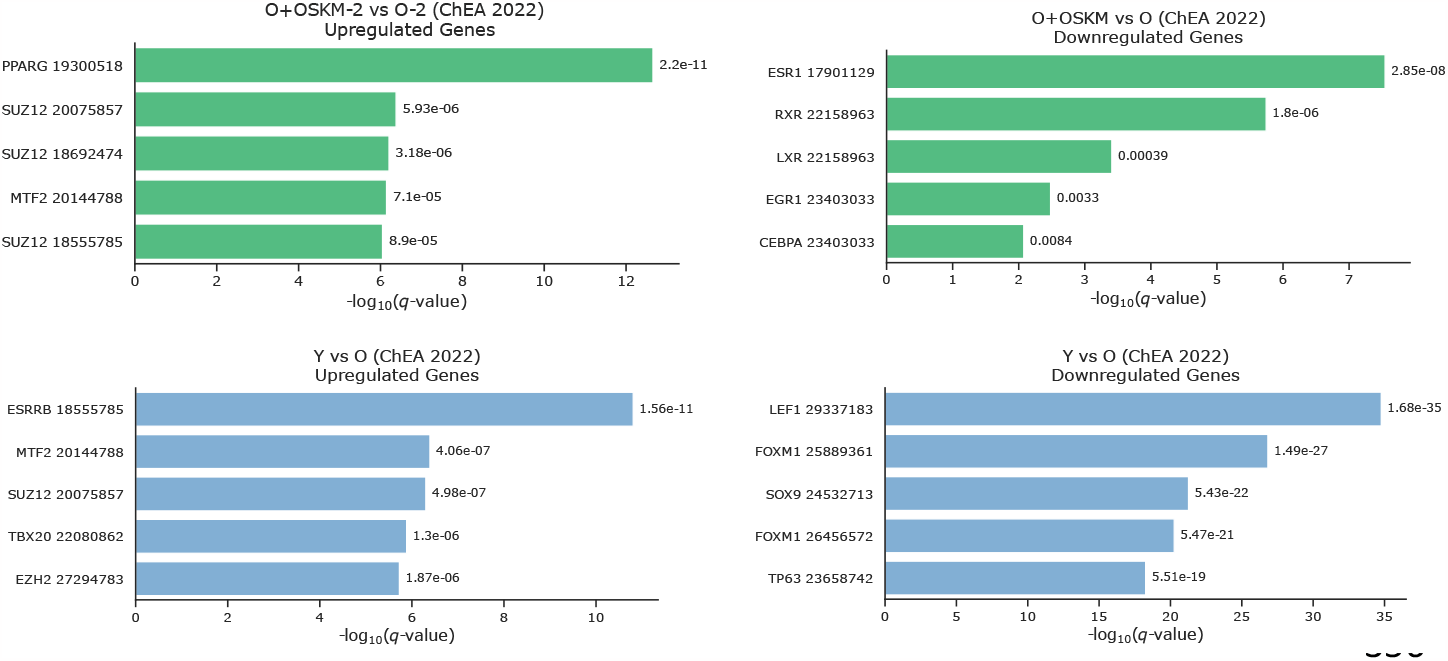
PRC2 targets are differentially expressed in aging and partial reprogramming-mediated rejuvenation. Enriched gene sets obtained from GSEA of the differentially expressed genes in young and treated old (old+OSKM) with untreated old comparisons.

Taken together, our results point to a central and previously unappreciated role for PRC2 signaling in the regulation of tissue aging and rejuvenation. Previous studies have shown an increase in mean DNA methylation levels in PRC2 target regions associated with aging^12,13^. Here we report a novel analysis of the aging epigenome based on epigenetic discordance as captured by the JSD, identifying PRC2 domains as the most epigenetically disrupted regions during aging due to hypermethylation and significant gain of entropy. Most importantly, we show that long-term partial reprogramming treatment is able to reverse such epigenetic disruption, decreasing both the mean methylation and entropy levels. Entropy as well as mean levels of DNA methylation have been associated with specific DNA binding motifs and regulatory DNA involved in developmental plasticity^14^, indicating such measures can unveil more comprehensive information than measures of mean differences alone. We report relevant results related to biological aging as opposed to chronological aging because of the convergence of PRC2 targets among both differentially methylated and differentially expressed genes, and the relevance of some of the most epigenetically altered genes to biological aging and regeneration.

## METHODS

### Animal use and care

All animal procedures were performed according to National Institutes of Health (NIH) guidelines and approved by the Committee on Animal Care at the Salk Institute.

Mice carrying the OSKM polycistronic cassette and rtTA were obtained from Jackson Laboratory (stock no. 011011). All the mice were in a C57BL/B6 background. The protocols for cyclic induction of OSKM and skin sample collection are described previously^5^.

### WGBS library preparation and sequencing

Genomic DNA was isolated using the MasterPure DNA Purification kit (Epicentre). We confirmed integrity of genomic DNA by gel electrophoresis. WGBS single indexed libraries were generated using NEBNext Ultra DNA library Prep kit for Illumina (New England BioLabs) according to the manufacturer’s instructions with the following modifications: 500 ng input gDNA was quantified by Qubit dsDNA BR assay (Invitrogen) and spiked with 1% unmethylated Lambda DNA (Promega, cat # D1521) to monitor bisulfite conversion efficiency. We fragmented input gDNA by Covaris S220 Focused-ultrasonicator to an average insert size of 350 bp. Samples were sheared for 60 sec using Covaris microTUBEs, with instrument settings of duty cycle 10%, intensity 5 and cycles per burst 200. Size selection was performed using AMPure XP beads and insert sizes of 300-400 bp were isolated. Samples were bisulfite converted after size selection using EZ DNA Methylation-Gold Kit or EZ DNA Methylation-Lightning Kit (Zymo cat#D5005, cat#D5030) following the manufacturer’s instructions. After bisulfite conversion, we performed amplification using Kapa Hifi Uracil+ (Kapa Biosystems, cat# KK282) polymerase based on the following cycling conditions: 98°C 45s / 8cycles: 98°C 15s, 65°C 30s, 72°C 30s / 72°C 1 min.

AMPure cleaned-up libraries were run on the 2100 Bioanalyzer (Agilent) High-Sensitivity DNA assay and samples were also run on the Bioanalyzer after shearing and size selection for quality control purposes. We quantified libraries by qPCR using the Library Quantification Kit for Illumina sequencing platforms (Kapa Biosystems, cat#KK4824) and the 7900HT Real Time PCR System (Applied Biosystems). We sequenced WGBS libraries on an Illumina HiSeq4000 instrument using 150 bp paired-end indexed reads and 25% of non-indexed PhiX library control (Illumina). We indicate coverage and average GpG depth in Table S2. The bisulfite conversion rate of unmethylated Lambda DNA was 99.5% on average.

### Mapping and quality control of whole genome bisulfite sequencing

FASTQ files were processed using Trim Galore! v.0.4.0 (Babraham Institute) to perform single-pass adapter- and quality-trimming of reads. FastQC v.0.11.2 was employed for quality control of reads. Reads were aligned to the mm10 genome using Bismark v.01.14.5. Separate M-bias plots for read 1 and read 2 were generated by running the Bismark methylation extractor using the ‘mbias_only’ flag, and these plots were used to determine how many bases to remove from the 5′ end of reads. The number was generally higher for read 2, which is known to exhibit a lower quality. The amount of 5′ trimming ranged from 5 bp to 20 bp. BAM files were subsequently processed with Samtools v.0.1.19 for sorting, merging, duplicate removal and indexing.

### Genomic features and annotations

Files and tracks bear genomic coordinates for mm10. We obtained CGIs annotatons from the UCSD Genome Browser. We defined CGI shores as sequences flanking 2-kb on either side of CGIs, shelves as sequences flanking 2-kb beyond the shores, and open seas as everything else. We used the R package ‘TxDb.Mmusculus.UCSC.mm10.knownGene’ to define genes, exons, and introns. We defined the promoter region of a gene as the 4-kb window centered at its transcription start site (TSS) and determined the gene body region to be the remainder of the gene. ChIP-seq and chromHMM data from mouse embryonic stem cells were obtained from the Mouse Encode Project.

### PEL computation

We computed DNA methylation potential energy landscapes (PELs) from WGBS data using informME (v0.3.2), a freely available information-theoretic pipeline for methylation analysis based on the 1D Ising model of statistical physics. For these computations, the entire genome was partitioned into consecutive non-overlapping genomic windows of 3-kb each and a PEL was estimated within each window from available WGBS reads using a maximum-likelihood approach^8,9.^

### Methylation level and entropy

Probability distributions of methylation levels, as well as mean methylation levels and normalized methylation entropies, were computed within each analysis region using informME^9^. The mean methylation level is the expected value of the methylation level *L* within an analysis region, and is given by 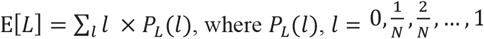, is the associated probability distribution of *L*. The normalized methylation entropy is a normalized version of Shannon’s entropy, given by *h* = −[1/log_2_(*N +* 1)] ∑_*l*_ *P*_*L*_ (*l*) log_2_ *P*_*L*_ (*l*), and was used to quantify the amount of methylation stochasticity observed within an analysis region. It ranges between 0 and 1, taking its maximum value when all methylation levels within an analysis region are equally likely (fully stochastic methylation), and achieving its minimum value only when a single methylation level is observed (perfectly ordered methylation).

### Jensen-Shannon distance and mutual information

Within an analysis region, the Jensen-Shannon distance between the two probability distributions 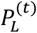 and 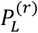 of the methylation level in test (all young/old treated) samples and a reference (Old+OSKM-2) sample was calculated by 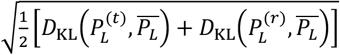, where 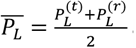 and 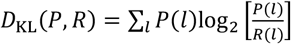 is the Kullback-Leibler divergence between two probability distributions *P* and *R* (also known as the relative entropy), and was used to quantify dissimilarities between the two probability distributions 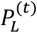 and 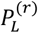. It ranges between 0 and 1, taking its minimum value only when the two probability distributions are identical, in which case no statistical discordance in methylation level is present, and its maximum value of 1 only when the supports of the two probability distributions do not intersect each other, in which case a maximum statistical discordance in methylation level is observed.

Within an analysis region, the mutual information between the methylation level *L* and the phenotype *A* is given by 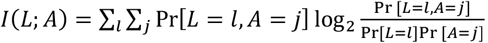, where *j* = 0 for the reference phenotype and *j* = 1 for the test phenotype, respectively. We used this quantity to measure, within an analysis region, the degree of mutual dependence between the methylation level and the phenotype, with higher values indicating a stronger mutual dependence. By making the (reasonable) assumption that the test and reference phenotypes are equally probable (i.e., Pr[*A*= 0] = Pr [*A*= 1] = 1/2), it can be shown that the average mutual information between the methylation level and the phenotype within a region of the genome that includes *K* analysis regions, given by 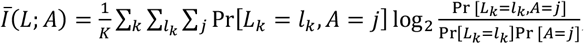, where *L*_*k*_ is the methylation level of the *k*-th analysis region, equals the square of the magnitude 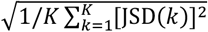 of the Jensen-Shannon distance within the genomic region, where JSD(*k*) is the Jensen-Shannon distance within the *k*-th analysis region. Note that the average mutual information ranges between 0 and 1, with higher values indicating a more informative genomic region in which there is a greater degree of mutual dependence between the methylation level and the phenotype. Notably, the previous relationship implies that the mutual information between the methylation level and the phenotype within an analysis region equals the square of the Jensen-Shannon distance between the probability distributions of the methylation level within the region in a test/reference comparison.

### Differential methylation analysis

We performed differential analysis between test (young or old treated) and reference (old untreated) on WGBS samples using informME^9^. We computed, within analysis regions, Jensen-Shannon distances (JSDs) between the corresponding methylation level probability distributions, as well as differences between mean methylation levels (dMMLs) and normalized methylation entropies (dNMEs).

### Hypothesis testing and gene ranking

In a single test/reference comparison, we computed within the promoter and body regions of each gene in the genome the magnitude of the Jensen-Shannon distance (JSD), which we calculated as the square root of the average of the squared JSD values within all analysis regions that overlap each feature (promoter or body). By following a previous statistical methodology^10^, we performed hypothesis testing to test against the null hypothesis that the JSD magnitude within a particular genomic feature (promoter or body) can be explained by normal technical, statistical, or biological variability. We did so by empirically constructing a null distribution for the values of all JSD magnitudes genome-wide, which we obtained by comparing our three young 4F samples. To account for variability in the number of analysis regions overlapping each genomic feature, we employed generalized additive models for location scale and shape (GAMLSS) with a logit skewed Student’s *t*-distribution. We also performed hypothesis testing simultaneously for methylation discordance within a gene’s promoter and body using Fisher’s summary statistic to test the null hypothesis that epigenetic discordance observed within a gene’s promoter or body in a test/reference comparison is only associated with biological, statistical, or technical variability in the reference samples, against the alternative hypothesis that this discordance is due to other factors within at least one of the two features considered (promoter or body). To evaluate genes in multiple test/reference comparisons, we used Fisher’s summary test statistic to test the null hypothesis that epigenetic discordance observed within a gene’s promoter and body in the test/reference comparisons is only associated with biological, statistical, or technical variability in the reference samples, against the alternative hypothesis that this discordance is due to other factors within at least one of the two genomic features considered (promoter or body) in at least one of the test/reference comparisons, and followed a similar approach to evaluate genes using only their promoters or bodies. We finally scored each gene by using the computed *p*-value for rejecting the null hypothesis and produced a ranked list of genes with increasing *p*-values, breaking possible ties by combining the *p*-value rankings obtained from each single test/reference comparison using the method of rank products. Finally, we evaluated the statistical significance of each ranking while controlling for the false-discovery rate (FDR) at 0.05 using *q*-values computed by the Benjamini-Hochberg (BH) procedure.

### Differentially gene expression analysis

RNA-seq reads were mapped to the mm10 genome and transcript-level quantification was performed using Salmon v.1.9.0. We used tximport v.1.2.0 to compute normalized gene-level counts from the transcript-level abundance estimates (scaling these using the average transcript length over samples and the library size). Only genes with at least 1 cpm in at least 3 samples were retained for downstream analysis. Differential gene expression was calculated using DESeq2 v.3.15, with trimmed mean of M-values normalization and multiple hypothesis correction of p-values performed using the Benjamini–Hochberg method. We tested for differential expression of genes in two comparisons: (1) young vs untreated old and (2) treated old vs untreated old. For a gene to be called a differentially expressed gene (DEG) it had to have a Benjamini-Hochberg adjusted P-value < 0.05 with no minimum log2 fold change cutoff.

### Gene Set Enrichment Analysis (GSEA)

GSEA was performed using two databases of gene sets; (1) Chromatin Enrichment Analysis (ChEA) 2022 and (2) Molecular Signatures Database (MSigDB) v.7.0.

## ACKNOWLEDGEMENTS

This work was supported by the Bloomberg Philanthropies (A.F.), Johns Hopkins University funding of the XDBio graduate program (to O.C.). The funders had no role in study design, data collection and analysis, decision to publish, or preparation of the manuscript.

