## Supplementary Figures for "Convergence of aging- and rejuvenation-related epigenetic alterations on PRC2 targets"

### Supplementary Materials

**Supplementary Figure 1. Distributions of MML and NME within selected genomic regions**

Boxplots showing the distributions of MML and NME values in young, treated old (old+OSKM) and untreated old within different genomic regions.

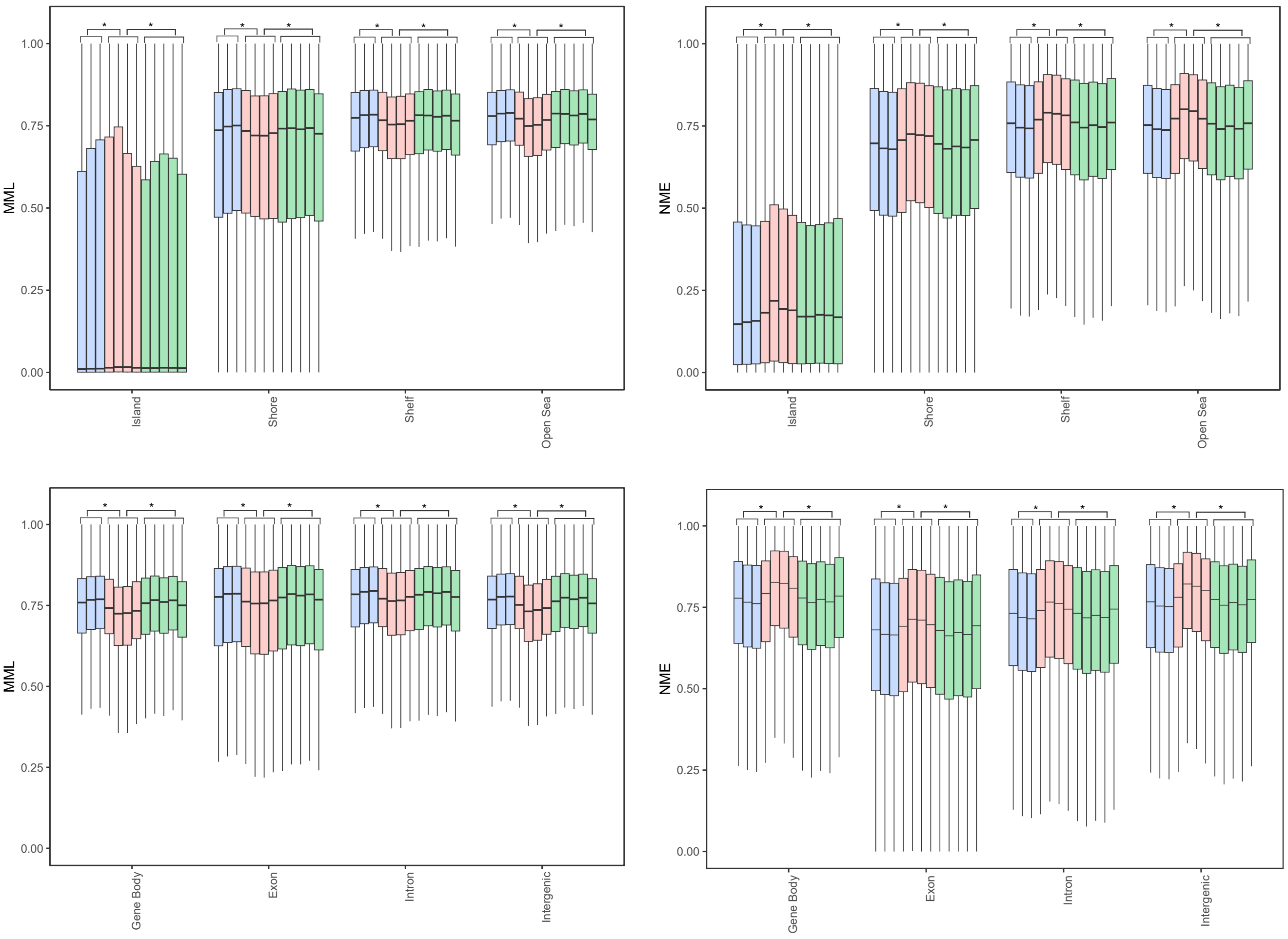

**Supplementary Figure 2. Distributions of JSD and differences in MML and NME across different chromatin states**

Boxplots showing the distributions of JSD and differences in MML (dMML) and NME (dNME) within different chromatin states in comparisons of representative treated old (O+OSKM-2) and young (Y-2) samples with untreated old (O-2) samples.

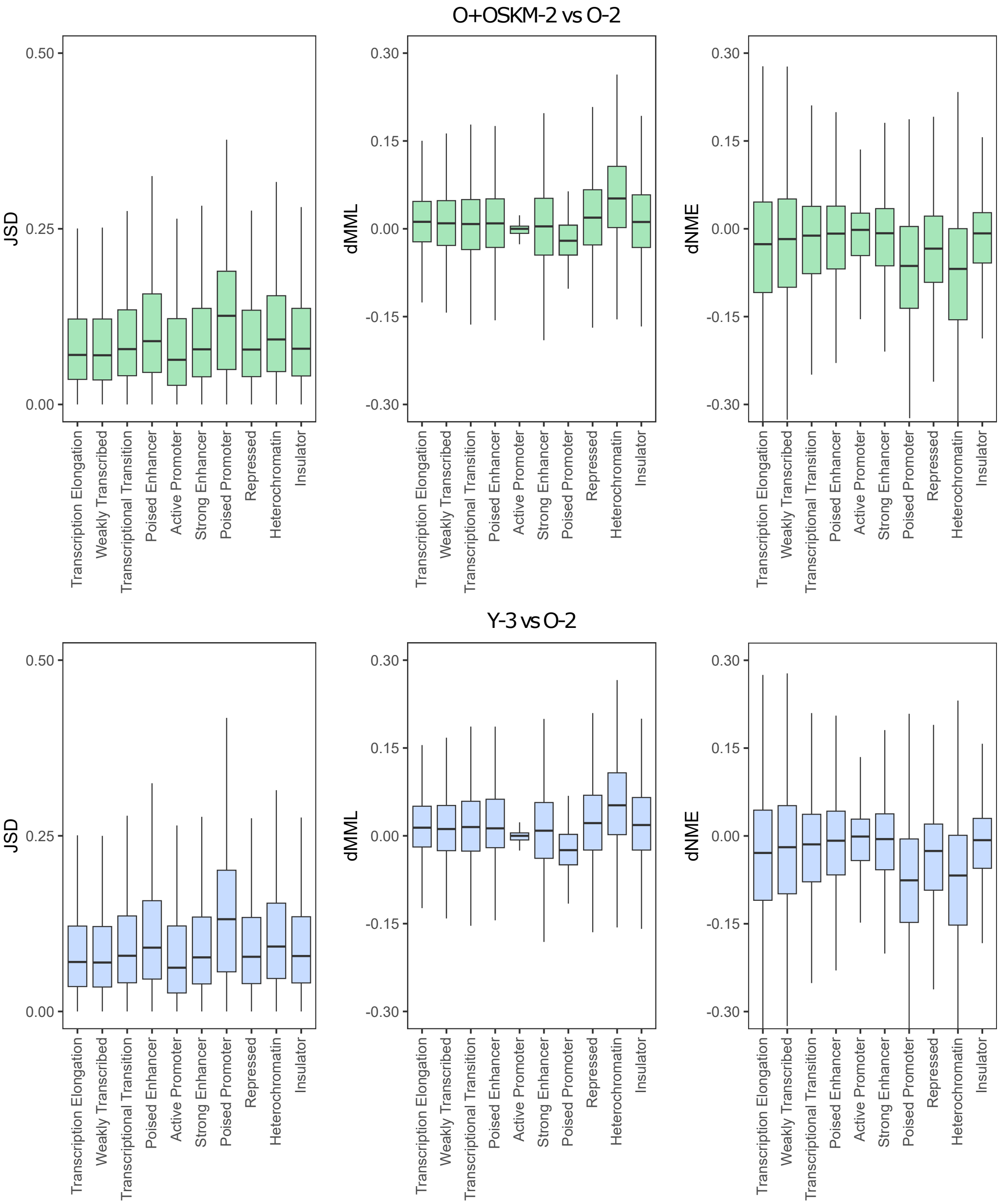

#### Supplementary Tables

**Supplementary Table S1.** Metadata for WGBS and RNA-seq samples

**Supplementary Table S2.** Ranked genes by epigenetic discordance between samples using Jensen-Shannon distance (JSD). Old+OSKM vs Old comparison

**Supplementary Table S3.** Ranked genes by epigenetic discordance between samples using Jensen-Shannon distance (JSD). Young vs Old comparison

**Supplementary Table S4.** Differential expression analysis. Old+OSKM vs Old comparison

**Supplementary Table S5.** Differential expression analysis. Young vs Old comparison
